# Travelling waves observed in MEG data can be explained by two discrete sources

**DOI:** 10.1101/2022.09.28.509870

**Authors:** Alexander Zhigalov, Ole Jensen

**Author notes:** Postal address: Centre for Systems Modelling and Quantitative Biomedicine, Institute of Metabolism and Systems Research, IBR Tower, Level 3, College of Medical and Dental Sciences, Birmingham, B15 2TT.

## Abstract

Growing evidence suggests that travelling waves are functionally relevant for cognitive operations in the brain. Several electroencephalography (EEG) studies report on a perceptual alpha-echo, representing the brain response to a random visual flicker, propagating as a travelling wave across the cortical surface. In this study, we ask if the propagating activity of the alpha-echo is best explained by a set of discrete sources mixing at the sensor level rather than a cortical travelling wave. To this end, we presented participants with gratings modulated by random noise and simultaneously acquired the ongoing MEG. The perceptual alpha-echo was estimated using the temporal response function linking the visual input to the brain response. At the group level, we observed a spatial decay of the amplitude of the alpha-echo with respect to the sensor where the alpha-echo was the largest. Importantly, the propagation latencies consistently increased with the distance. Interestingly, the propagation of the alpha-echoes was predominantly centro-lateral, while EEG studies reported mainly posterior-frontal propagation. Moreover, the propagation speed of the alpha-echoes derived from the MEG data was around 10 m/s, which is higher compared to the 2 m/s reported in EEG studies. Using source modelling, we found an early component in the primary visual cortex and a phase-lagged late component in the parietal cortex, which may underlie the travelling alpha-echoes at the sensor level. We then simulated the alpha-echoes using realistic EEG and MEG forward models by placing two sources in the parietal and occipital cortices in accordance with our empirical findings. The two-source model could account for both the direction and speed of the observed alpha-echoes in the EEG and MEG data. Our results demonstrate that the propagation of the perceptual echoes observed in EEG and MEG data can be explained by two sources mixing at the scalp level equally well as by a cortical travelling wave. This conclusion however does not put into question continuous travelling waves reported in intracranial recordings.

## Introduction

Cortical travelling waves has been proposed as a potential mechanism supporting communication and computations in the brain (Das et al., 2022; Visser et al., 2017; Zhang et al., 2018). Mathematical models further explain the functional advantages of the travelling waves in the brain (Ermentrout & Kleinfeld, 2001; Muller et al., 2018), and importantly, the case has been made that neocortical neurons can generate travelling waves (Bhattacharya et al., 2022) that can be detected with EEG (Nunez, 1974).

Travelling waves have been identified using both invasive and non-invasive EEG recordings. While invasive recordings provide a high signal-to-noise ratio, the intracortical electrodes cover a relatively small area of the brain. In contrast, non-invasive EEG provides whole-head coverage, but the signals are strongly affected by spatial mixing and noise. These discrepancies between the techniques introduce inconsistency across studies. First, the reported velocity of the travelling waves varies within a wide range, 0.1 – 10 m/s (Alamia & VanRullen, 2019), suggesting different origins of the waves. Second, there is inconsistency in the direction of the travelling waves. While intracortical recordings demonstrate different directions of propagation within relatively small areas in the prefrontal and temporal cortices (Bahramisharif et al., 2013; Davis et al., 2020; Dickey et al., 2021; Halgren et al., 2019), non-invasive EEG studies report large-scale propagation predominantly in occipital-to-frontal direction (Alamia & VanRullen, 2019; Lozano-Soldevilla & VanRullen, 2019; Pang et al., 2020). Third, EEG systems require a reference sensor, which might also impact the source modelling of the travelling waves. This problem can be partially alleviated by using MEG which provides whole-head coverage and does not require a reference sensor.

Many studies have reported travelling waves associated with endogenous brain rhythms that are not dependent on external stimulation (Benucci et al., 2007; Muller et al., 2018; Sato et al., 2012). However, broad-band visual flicker may also be used to evoke responses, so-called perceptual echoes, that are organised as travelling waves (Alamia & VanRullen, 2019; Lozano-Soldevilla & VanRullen, 2019). The propagation of the perceptual echoes in the 8-13 Hz alpha band has so far been studied using only EEG, and thus, MEG may provide new insight into the origin of the travelling alpha echoes due to its superior spatial resolution. Importantly, previous EEG studies have highlighted the possibility that travelling waves at the scalp level may not only be explained by travelling cortical activity but could also be a result of a linear mixture of neuronal activity generated by two phase-lagged dipoles (Lozano-Soldevilla & VanRullen, 2019) which can be tested using MEG.

In this study, we applied broadband visual stimulation to participants while we recorded the ongoing MEG. The perceptual echoes were estimated from the temporal response functions derived from the visual broadband input and the MEG sensors. We further applied source modelling as well as forward simulation to understand the neuronal sources explaining the propagation of the perceptual alpha-echoes.

## Methods

### Participants

Twenty-four participants (mean age: 33; age range: 18-38; 14 females) with no history of neurological disorders partook in the study. Six participants were excluded from the analysis because the perceptual echo was at the noise level. The study was approved by the local ethics committee (University of Birmingham, UK) and written informed consent was acquired before enrolment in the study. All participants conformed to standard inclusion criteria for MEG experiments. Participants had normal or corrected-to-normal vision.

### Experimental paradigm

Two grating stimuli, 5.8 degrees of visual angle, were presented bilaterally on a projector screen, placed at 1.4 meters from the participants’ eyes (Fig. 1A). After 1 s from the stimuli onset, the left and right gratings started contracting for 3 s either coherently (same direction) or incoherently (different directions). The direction of the motion (up / down) of the left and right grating stimuli was random in consequent trials. The grating stimuli moved at a constant speed of 0.5 degree/s. The participants were instructed to focus on the fixation point and press the button when a cue indicating the direction of motion (i.e. coherent or incoherent) occurred at the fixation point. If the direction of motion, either coherent or incoherent, matched the cue, equal or unequal sign, respectively, the participant was instructed to press the button with the middle finger; otherwise with the index finger. The response instructions, i.e. left or right hand, were counterbalanced across the participants.

**Fig. 1.**
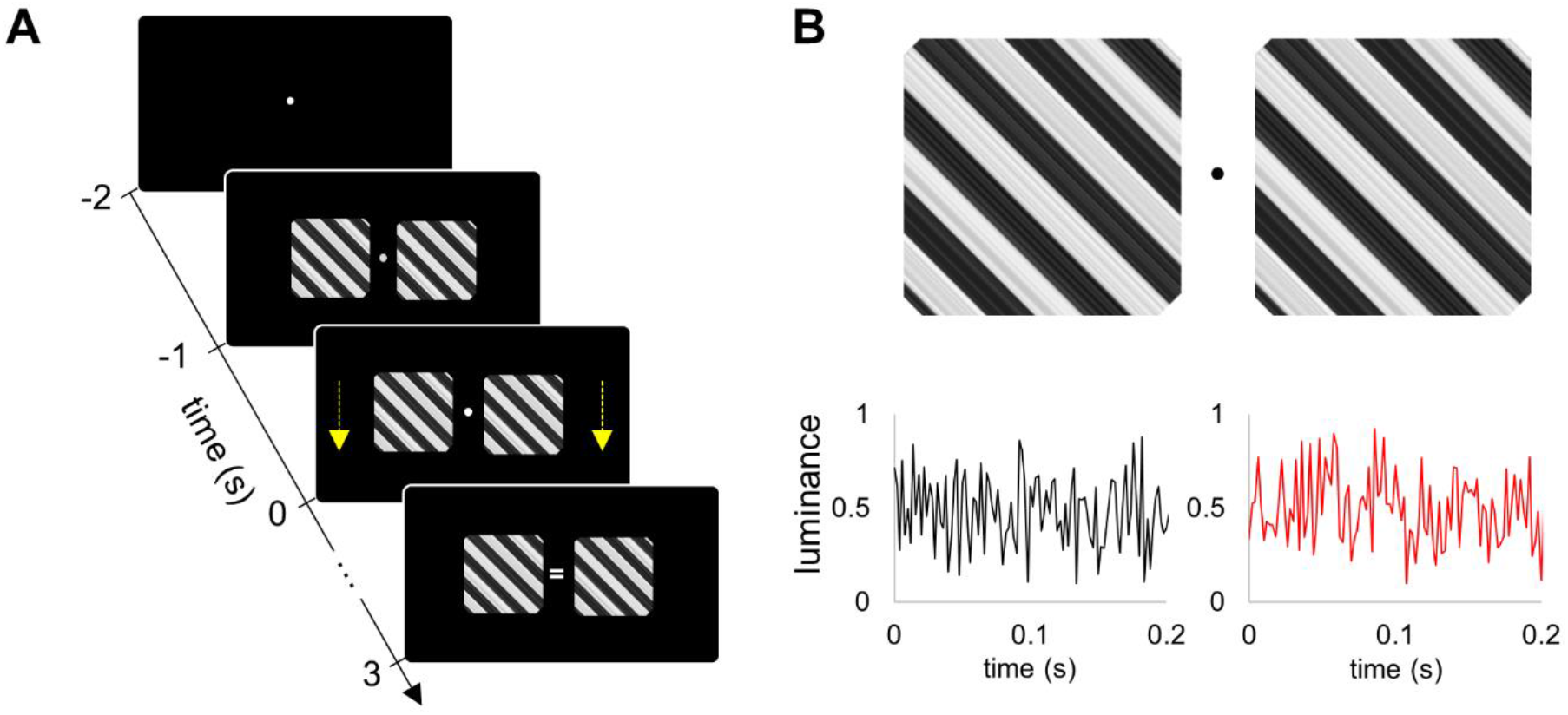
Experimental paradigm. A) Two grating stimuli with either coherent or incoherent motion were presented bilaterally. Motion started 1 s after the onset of the flickering stimuli and lasted for 3 s. (B) The luminance of the left and right grating stimuli was modulated by two independent broadband signals.

The luminance of the stimuli was modulated using a broadband (i.e. noise with uniform distribution) flickering signal (Fig. 1B). The broadband signals for the left and right stimuli were uncorrelated (the correlation was below 0.01). We used the PROPixx DLP LED projector (VPixx Technologies Inc., Canada) to present the visual stimuli at a high refresh rate of 1440 Hz with a resolution of 960 x 600 pixels [see, (Zhigalov & Jensen, 2020)]. The experimental paradigm was implemented in MATLAB (Mathworks Inc., Natick, USA) using Psychophysics Toolbox 3.0.11 (Pelli, 1997). (Pelli, 1997).

### MEG and MRI acquisition

Prior to the measurement, the head position of the participant was digitized using the Polhemus Fastrack electromagnetic digitiser system (Polhemus Inc., USA). The MEG data were acquired using a 306-sensor TRIUX Elekta Neuromag system (Elekta, Finland), while the participant was sitting in an upright position (60°) in a dim light magnetically shielded room. The magnetic signals were bandpass filtered within 0.1 – 330 Hz with embedded anti-aliasing filters and sampled at 1000 Hz. Simultaneously with the MEG, we also acquired the coordinates of the pupil and the pupil diameter using an EyeLink eye-tracker (SR Research, Canada).

The broadband flickering (or tagging) signals were recorded using a custom-made photodetector (Aalto NeuroImaging Centre, Aalto University, Finland) that was connected to a miscellaneous channel of the MEG system. This allowed us to acquire the tagging signal with the same temporal precision as the MEG data.

Additionally, a high-resolution T1-weighted anatomical image (TR/TE of 7.4/3.5 ms, a flip angle of 7°, FOV of 256×256×176 mm, 176 sagittal slices, and a voxel size of 1×1×1 mm3) was acquired using 3-Tesla Phillips Achieva scanner.

### MEG data preprocessing

Since magnetometer signals are strongly affected by the noise, additional pre-processing methods are needed to attenuate external interference and sensor artifacts. To this end, we used MNE-Python tools (Gramfort et al., 2013). MNE-Python package allowed performing signal-space separation (SSS; (Taulu & Simola, 2006)) and Maxwell filtering together. While SSS decomposes MEG signal into components with sources inside and outside the measurement volume and discards the external components related to environmental noise, Maxwell filtering omits the higher-order components of the internal subspace, which are dominated by the sensor noise.

To further suppress artifacts caused by the cardiac signal and blinks, we applied independent component analysis [ICA; (Hyvarinen, 1999)]. Prior to the analysis, the MEG time series were bandpass filtered (1 – 99 Hz) and down-sampled to 250 Hz. The components with topographies corresponding to blinks and cardiac signals were projected out as follows (Oostenveld et al., 2011),

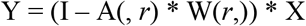

where X represents the original MEG data; A and W are mixing and unmixing matrices respectively; I is an identity matrix; r denotes indices of the components corresponding to blinks and cardiac signal; Y represents MEG data with projected out artifacts.

The artifact-free MEG time series were segmented into 6 s epochs; -2 to 4 s relative to the onset of the stimulus motion. Although ICA allowed removing blinks from the MEG signal, we additionally analysed the X-axis and Y-axis channels of the eye-tracker, to exclude trials containing blinks and saccades. The eye blinks were detected in the eye-tracker channels by z-scoring the time series and applying a threshold of 5 SD was applied so that any deflection above the threshold was classified as a blink. The saccades were detected using a scatter diagram of X-axis and Y-axis time series of the eye-tracker for each trial. An event was classified as a saccade if the gaze shifted away from the fixation point by 2° and its duration was more than 500 ms. Trials contaminated by blinks and saccades were removed from further analysis. We also rejected trials containing large-amplitude events (above 5 SD) in MEG which are mainly associated with motion and muscle artefacts. As a result, the number of trials that remained after exclusion was 304 ± 12 (mean ± SD) per participant. For each participant, the number of trials per condition was equalized by randomly selecting the same number of trials.

### Temporal response functions (TRF)

Prior to computing TRF, the magnetometer signals were bandpass filtered within 1 – 99 Hz.

The TRFs were computed between the broadband flickering signal and the MEG gradiometers for each trial. To this end, we applied the cross-correlation approach (VanRullen & Macdonald, 2012),

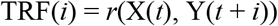

where X and Y denote input tagging signal and MEG, respectively; *i* is the time lag ranging from - 0.25 to 0.75 s with a step of 0.001 s; *t* is the vector of length 1.5 seconds; *r* is Person correlation; TRF denotes response function.

It should be noted that this approach is essentially equivalent to more sophisticated approaches such as ridge regression (Crosse et al., 2016) in case if the tagging signals are temporally uncorrelated.

The TRFs computed for each trial were then averaged across the trials.

### Source-space analysis

The alpha-echoes were computed in the source space using the linearly constrained minimum variance (LCMV) beamformer (Van Veen et al., 1997), as implemented in Fieldtrip (Oostenveld et al., 2011). To this end, we first reconstructed time series in the source space by applying LCMV beamformer to MEG data and then estimated the alpha-echoes (see, above) for each source point.

To build a forward model, we first segmented the MRI images using Fieldtrip, and then manually aligned the MRI images to the head shape digitization points acquired with the Polhemus Fastrak. Finally, a single shell head model was prepared using surface spherical harmonics fitting the brain surface (Nolte, 2003).

The individual anatomy was wrapped to the standard MNI template using non-linear normalisation (Oostenveld et al., 2011). The template grid had a 10 mm resolution, resulting in 6804 source points for each participant.

The covariance matrix was estimated in the time interval [0, 2.5] s relatively to the onset of stimuli motion. Since, the SSS method (see, above) reduces the rank of data, we estimated the rank after SSS and initialise the regularisation parameter kappa using the estimated rank. We computed the unit-noise-gain or weight-normalized beamformer (Westner et al., 2022) to avoid bias towards deep sources.

### Simulations using MEG and EEG models

As suggested elsewhere (Lozano-Soldevilla & VanRullen, 2019; Orczyk et al., 2021), the travelling waves might be explained by the two-source model, meaning that the propagation effect at the scalp level may occur as a result of the summation of two waves generated by distant sources. To test this hypothesis, we used a realistic head model and simulated the time series of two dipoles located in the occipital and parietal cortices, based on the empirical findings. The simulations were performed for both MEG and EEG source models using “ft_dipolesimulation” function from the Fieldtrip toolbox (Oostenveld et al., 2011). Importantly, we used the radial orientation of the sources in both MEG and EEG simulations.

## Results

In this study, we presented participants with grating stimuli while modulating the luminance of the stimuli by random noise, and simultaneously acquiring MEG. Temporally uncorrelated flickering stimuli were shown in the left and right hemifield.

### Alpha-echo

The brain response to the randomly flickering stimuli was estimated using temporal response functions (TRFs). The TFR reflects the kernels best explaining the brain response when convolved with the input stimuli. The individual temporal responses were averaged across the participants for both the right (Fig. 2A) and left stimulation sides (Fig. 2B). The TRFs are dominated by a 10 Hz response and have been termed “perceptual alpha echoes” (VanRullen & Macdonald, 2012). The topographies of the TFRs at 200 ms showed a dipolar response and the strongest responses were observed in sensors over parietal areas.

**Fig 2.**
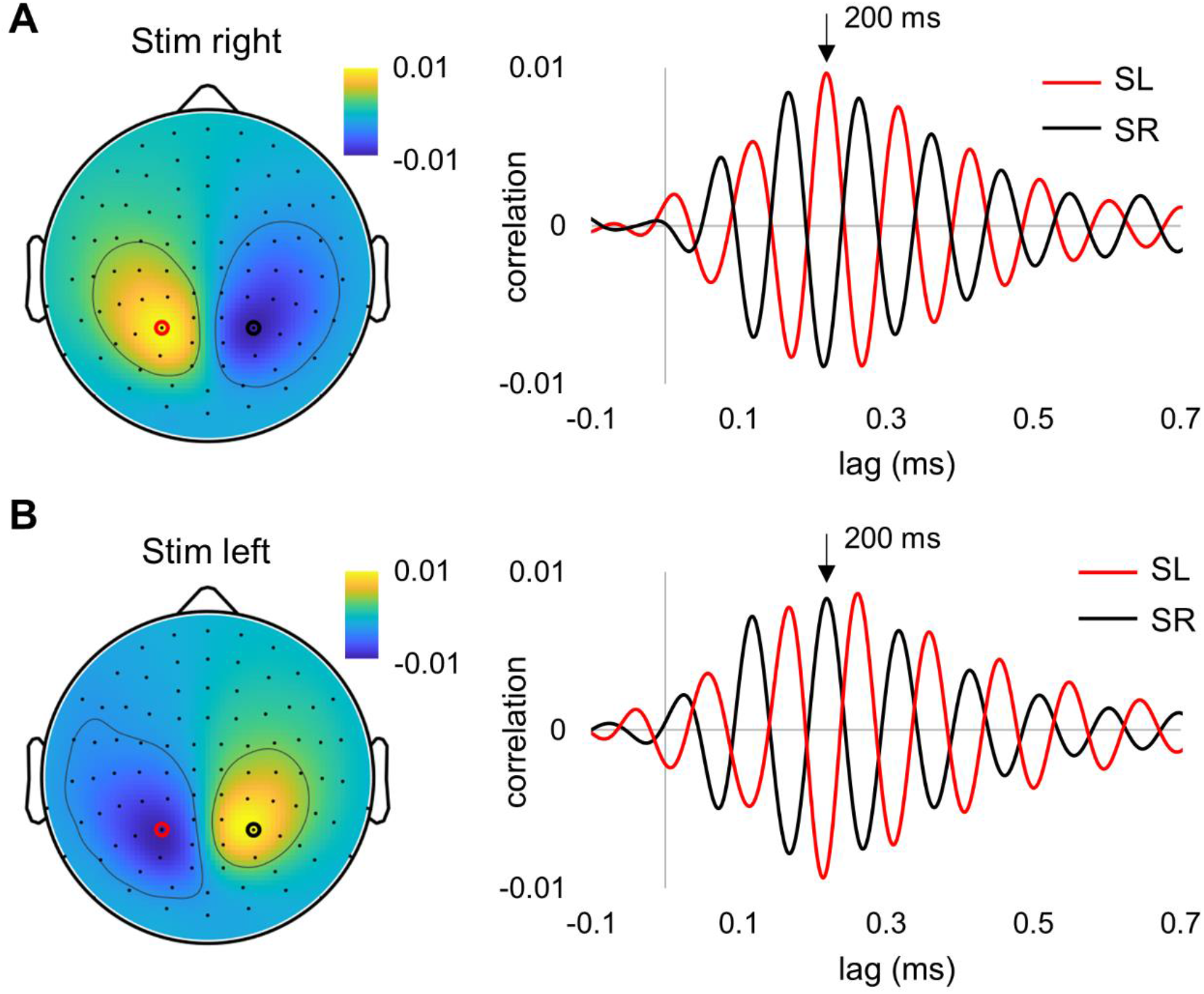
Group-level temporal response functions (TRFs) were computed for the right (A) and left (B) flickering signals. Magnetometers with the largest alpha-echo amplitude are depicted with black and red circles, and the corresponding alpha-echoes are shown on the right panel. The topography is shown for t = 200 ms.

### The alpha-echo in sensor and source spaces

Next, we considered the magnitude of the spatial distribution of the TRFs. Since the TRFs in the left and right hemispheres were in anti-phase, we considered the group-level responses contralateral to stimulation in the left and right hemispheres separately (Fig. 3A). We observed a systematic spatial decay of the TFR peak amplitude observed at around 200 ms, and the decay of the amplitudes with distance was nearly linear (*p* < 10^-12^). We then used LCMV beamformers to assess the power of TRFs in source space. To this end, the sensor-space MEG times series were projected into source space, and then the TRFs were computed for each dipole (or grid point). The maximum power of the TRF was localised in the parietal regions near the midline (along X-axis) for both the right and left stimulation sides (Fig. 3B).

**Fig 3.**
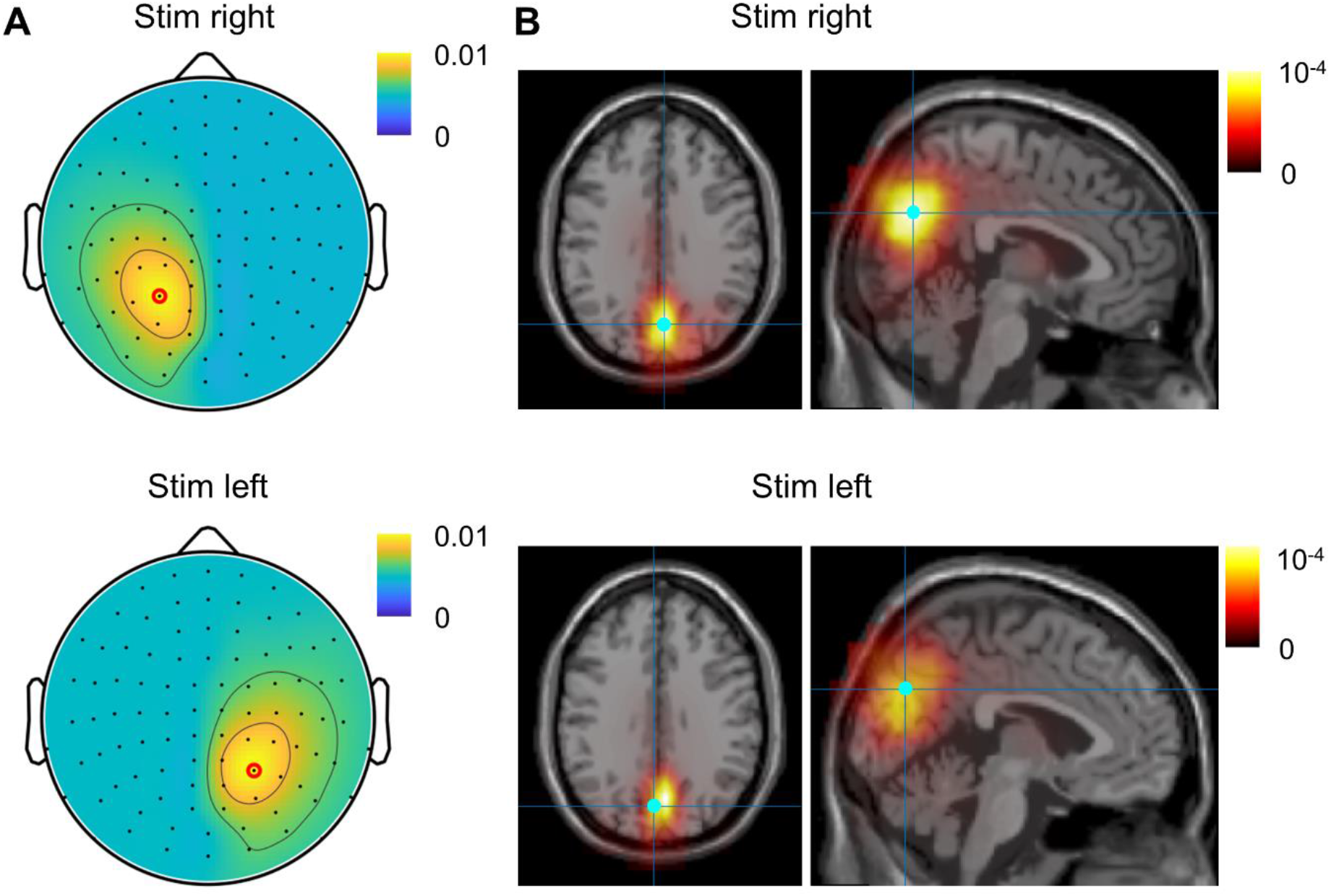
(A) The topography of the TFR amplitudes with respect to the right (top plot) and left (bottom plot) stimulation sides. (B) Distribution of the TRF power in source space with respect to the right (top plot) and left (bottom plot) stimulation sides. The voxel with the strongest response is indicated with blue dot.

### Propagation of the alpha-echo in sensor space

We next set out to quantify the propagation of the perceptual alpha echo in regard to the reference sensor with the strongest response. We did this by simply assessing the shift of the TRF in a neighbouring sensor with respect to the reference sensors. The latencies of the TRF relatively to the reference sensor showed a linear increase with distance (*p* < 10^-6^, *F*-test), predominantly in the centro-lateral direction (Fig. 4A). The topography revealed two clusters with respect to the left and right hemifield visual stimulation (Fig. 4A). The propagation velocity derived from the latencies was around 10 m/s which is higher compared to that reported using EEG (Alamia & VanRullen, 2019). Similarly to the sensor level, the TRF latencies in the source space showed two discrete clusters, rather than a continuous increase, in the parietal and occipital cortices (Fig. 4B). Note that we considered the latencies in a restricted area in the source space because signal-to-noise in the frontal regions was much lower compared to that in the parietal cortex where the response was strongest. Both sensor-and source-level results showed that the latencies of the TRF formed two clusters with similar values within the clusters, which does not support the idea of continuous propagation of the waves.

**Fig. 4.**
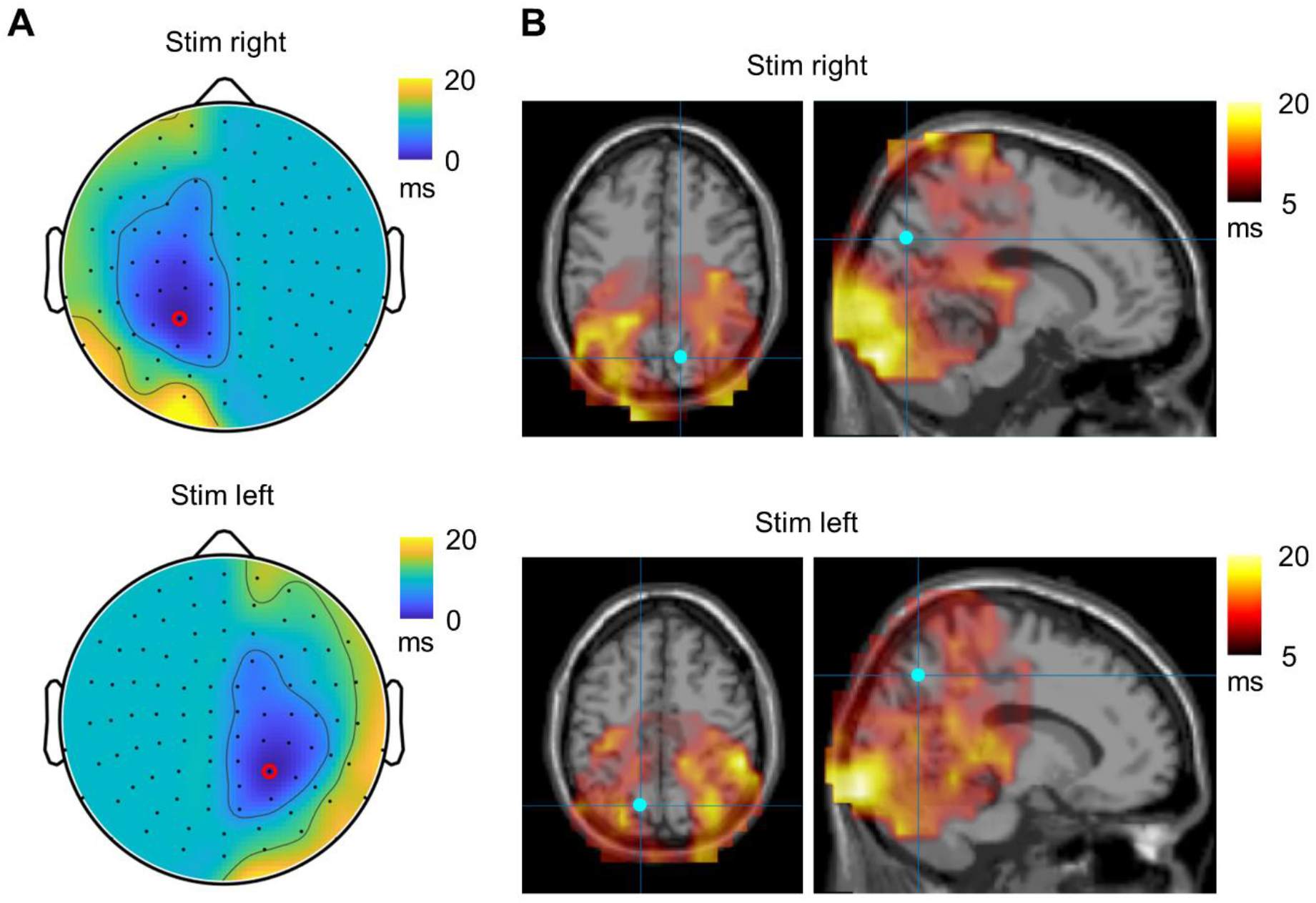
(A) The topography of the TFR latencies with respect to the right (top plot) and left (bottom plot) stimulation. The sensor with maximum TRF is indicated by a red circle. (B) Distribution of the TRF latencies in source space with respect to the right (top plot) and left (bottom plot) stimulation. The voxel with the largest TRF power is indicated by the blue dot.

### Spatiotemporal evolution of the sources

To obtain further insight into the spatiotemporal dynamics of the TRF, we evaluated the power of TRF in source space at each time lag. To this end, we searched for the voxels that showed the maximum power of TRF in the alpha band at each lag. The maximum power of the TRF peaked at around 100 and 200 ms, respectively (Fig. 5A). These peaks were associated with two voxels with highly stable coordinates (Fig. 5B). While the voxel associated with the first peak was located in the primary visual cortex (Fig. 5C), the voxel associated with the second peak was located in the parietal area (Fig. 5D). These results showed that the alpha-echo was clearly related to the second source with a peak in power at around 200 ms, and there is another source with a peak in power at around 100 ms located in the primary visual cortex. This observation may support an alternative hypothesis regarding the origin of travelling waves in the brain. Often, the travelling waves at the sensor level are explained by waves travelling over the brain surface and the fields are then projected to the sensors. Alternatively, as in our case, the travelling waves can be generated by two sources at different locations oscillating at different phases, and superposition of the respective fields at the sensors occurs as a travelling wave.

**Fig. 5.**
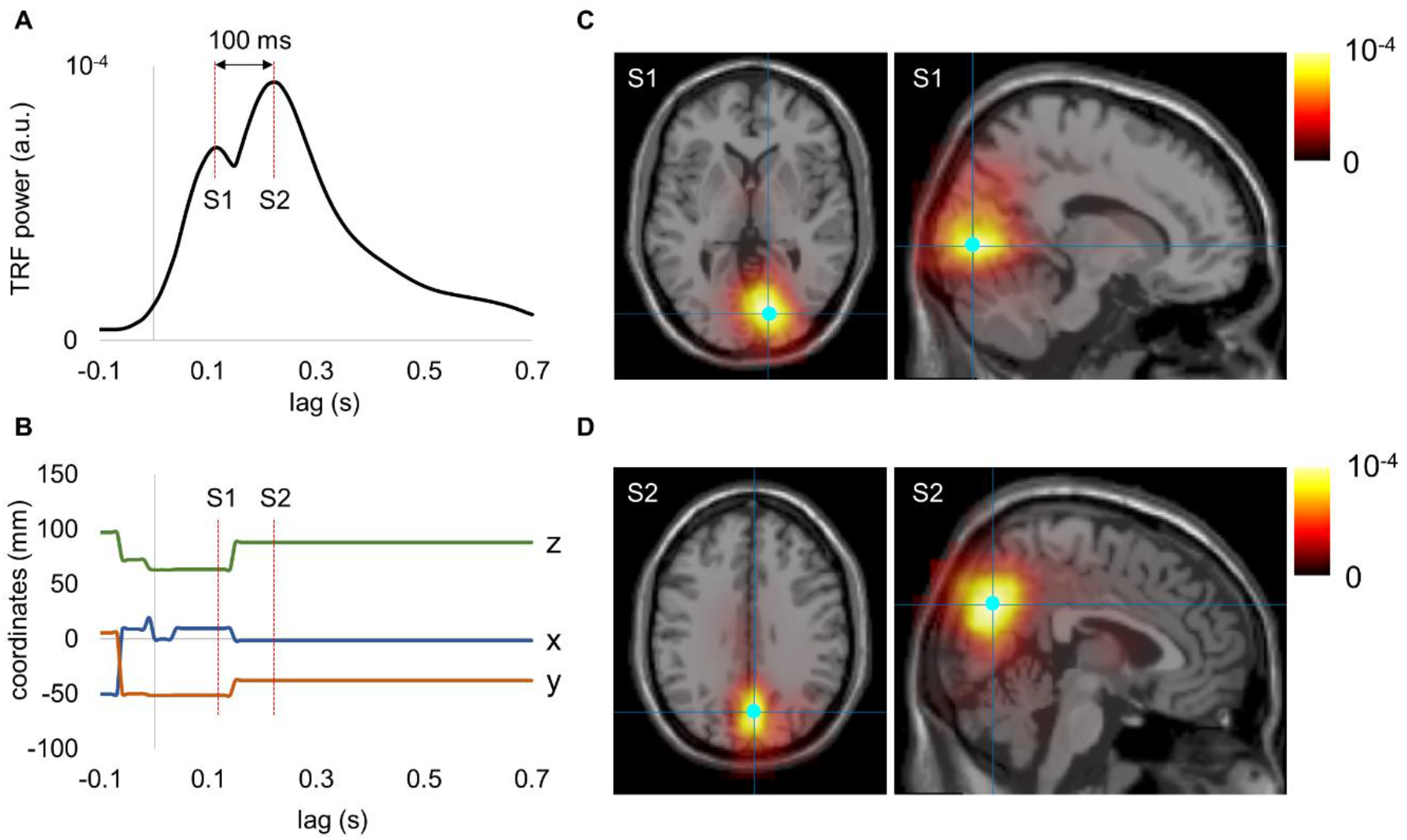
Spatiotemporal dynamics of the TRF power in the alpha band with stimulation on the left side. (A) The maximum power of TRF in the alpha band at each lag. (B) The coordinates of the voxel with maximum power in the alpha band at each lag. (C) The distribution of power associated with the first source (S1) peaked at around 100 ms. (D) The distribution of power associated with the second source (S2) peaked at around 200 ms.

### Simulations using MEG and EEG models

Using a realistic MEG forward model, we simulated the sensor’s level activity originating from two sources located in the occipital and parietal areas, in the right hemisphere, which were selected based on our empirical findings (Fig. 6A,B). The spatial patterns of the amplitudes and latencies of the simulated activity closely reproduced our empirical observations in MEG (Fig. 6C,D).

**Fig. 6.**
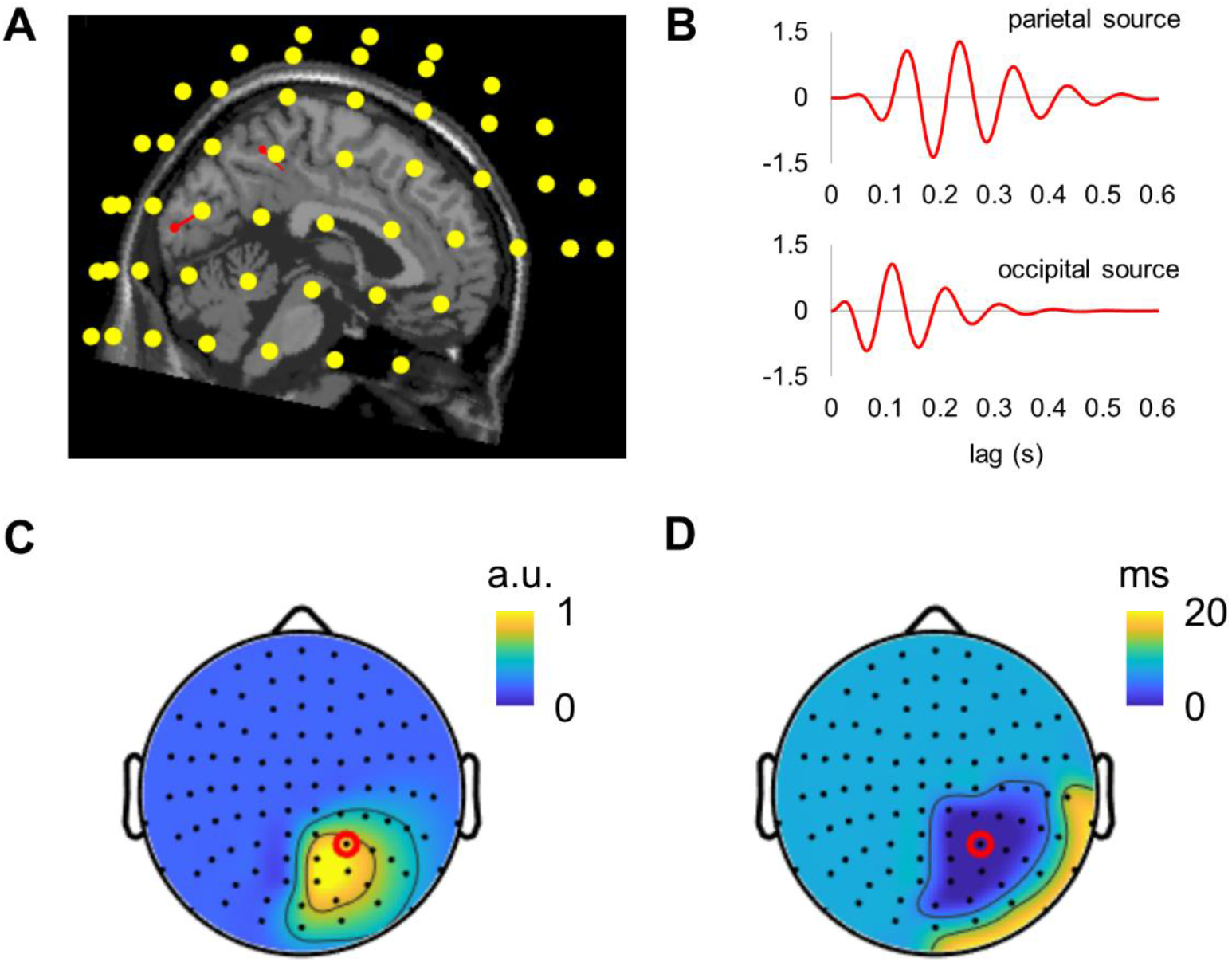
Simulated MEG activity from two sources closely resembles our empirical findings. (A) The location of the dipoles (red arrows) in the parietal and occipital regions relative to the MEG sensors. (B) Simulated activity associated with the parietal and occipital sources. (C) Topography of the amplitudes of the simulated alpha-echoes. (D) Topography of the latencies of the simulated alpha-echoes, relatively to the sensor with the strongest alpha-echo response (red dot).

To further relate our MEG findings to the results of previous EEG studies, we simulated EEG time series using the same dipole configuration as in our MEG simulations (Fig. 7A, B). The topography of the amplitude of perceptual echoes in EEG was similar to that in the MEG simulations (Fig. 7C). However, in contrast to MEG observations, the latencies of the echoes in simulated EEG showed a pronounced propagation pattern from occipital to frontal brain areas (Fig. 7D). Interestingly, the latencies between neighbouring sensors along the occipital-frontal axis showed nearly linear increase which resembled the results of previous EEG studies [e.g., (Alamia & VanRullen, 2019; Lozano-Soldevilla & VanRullen, 2019)]. These simulation results demonstrated that two dipoles located in the parietal and occipital cortices can produce the travelling waves at the sensor’s level, propagating from the occipital to the frontal areas.

**Fig. 7.**
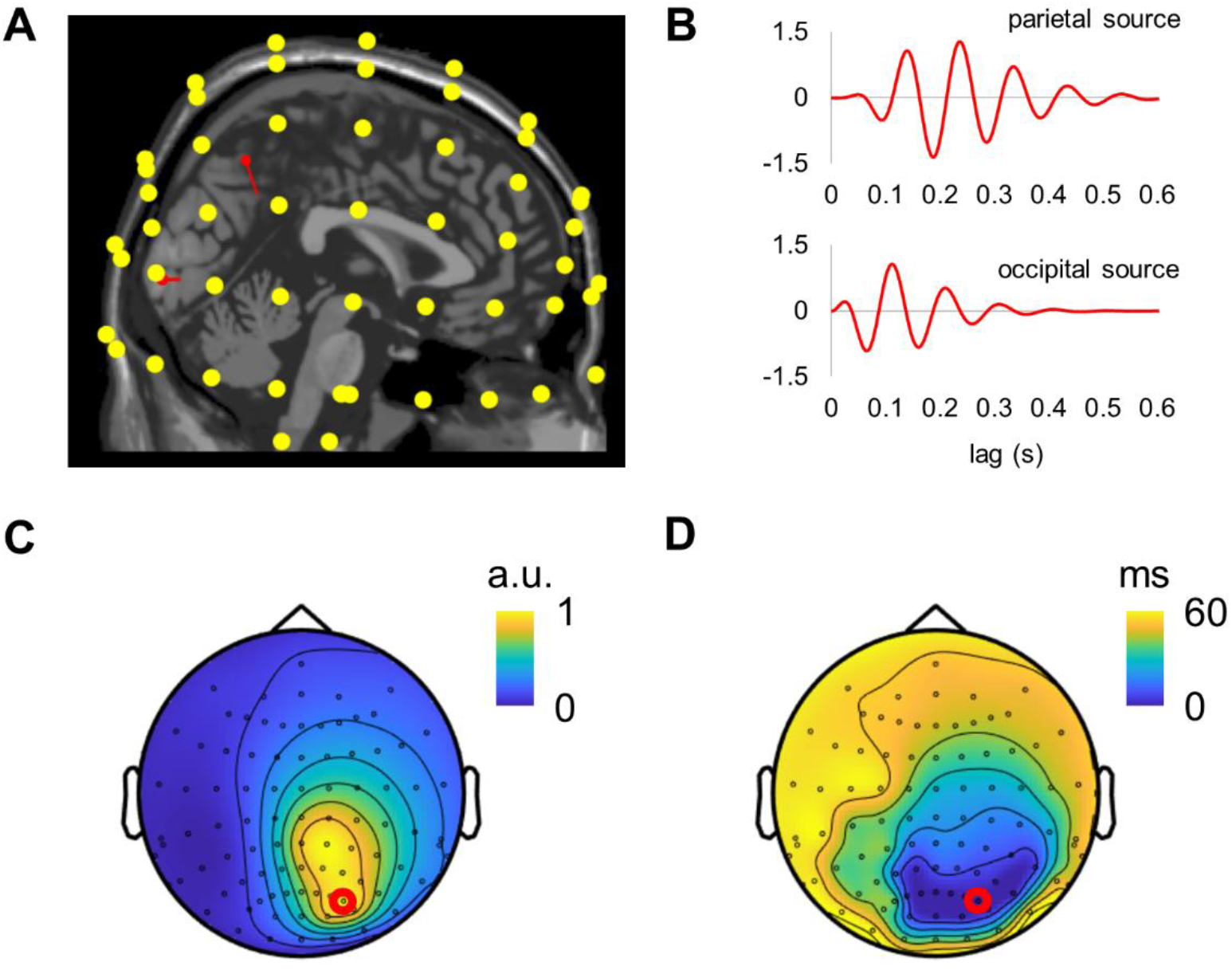
Travelling waves in the simulated EEG activity resemble those reported in previously EEG studies. (A) The location of the dipoles (red arrows) is relative to the EEG sensors. (B) Simulated activity associated with the parietal and occipital sources. (C) Topography of the amplitudes of the simulated alpha-echoes. (D) Topography of the latencies of the simulated alpha-echoes, relatively to the sensor with the strongest alpha-echo response (red dot).

## Discussion

In this study, we used broadband frequency tagging and MEG to assess the propagation of the alpha-echoes. Considering the better spatial resolution of MEG compared to EEG, we aimed at uncovering the neuronal sources explaining travelling waves in the brain. In contrast to previous EEG studies, we observed only a small (<20 ms) propagation delay of perceptual echoes across the scalp. Using source modelling, we found that the perceptual alpha-echoes were well explained by two generators in the primary visual and parietal cortices. This is consistent with the notion that travelling waves at the scalp level may occur as a result of a summation of neuronal activity generated by two phase-lagged dipoles. To further clarify this, we used a realistic forward model to estimate the activity generated by two sources, in occipital and parietal areas, in the MEG sensors. These stimulations explained the MEG findings well. Finally, we used an EEG forward model to project the two-dipole activity to EEG sensors. The sensor-level EEG travelling waves closely resembled those reported in previous EEG studies. Thus, our simulation results show that the sensor-level travelling waves reported in MEG and EEG can be explained by a common two-dipole model.

### Mechanism of travelling waves

Three mathematical models for generating travelling waves in the brain have been proposed (Ermentrout & Kleinfeld, 2001). These models are equally plausible and are built upon basic principles of neuronal coupling such as (*i*) delayed excitation from a single oscillator, (*ii*) propagating pulses in an excitable network, and (*iii*) phase-locked weakly coupled oscillators. Recently, Alamia and VanRullen (Alamia & VanRullen, 2019) proposed an further developed model to describe the propagation of the perceptual alpha band echoes in EEG. This model utilises a principle similar to propagating pulses in an excitable network and is composed of multiple hierarchically organised neuronal generators. Our results are consistent with such a model composed of two hierarchical levels, but not a multi-level model. Our source-space analysis confirmed the presence of two sources of perceptual alpha-echoes that were located in the primary visual and parietal cortices with peak amplitude at around 100 and 200 ms, respectively. The spatial and temporal dynamics of these sources essentially represent the stages of visual processing when activation in the primary visual cortex is followed by activation in the parietal cortex. Such two-level hierarchical processing can produce activity that appears as travelling wave at the scalp level.

### The direction of propagation of travelling waves

In contrast to EEG studies, our MEG results revealed a more focal propagation of the perceptual alpha-echoes, essentially restricted by the occipital and parietal areas. Interestingly, EEG studies reveal predominantly occipital-frontal propagation of the travelling waves for either a single flickering stimulus presented centrally (Pang et al., 2020) or two independent stimuli presented bilaterally (Lozano-Soldevilla & VanRullen, 2019). In contrast, the travelling waves in this MEG study demonstrated a tendency for centro-lateral propagation. Since the direction of propagation might be functionally relevant (e.g., (Aggarwal et al., 2022)), the discrepancy between MEG and EEG results makes it difficult to assess the functional relevance of the travelling waves. We conclude that sensor-level results should be considered with caution.

### Functional role of travelling waves

A growing body of evidence suggests that travelling waves are functionally relevant for memory and attention (Zhang et al., 2018). Although we have not observed a large-scale propagation of the perceptual echoes, we identified the sources and their dynamics, which can be further linked to cognitive functions, particularly, attention. Recently, we have shown that attention modulates the amplitude of the alpha-echo in the primary visual and parietal cortices (Lozano-Soldevilla & VanRullen, 2019) and that the alpha oscillations do not implement the gain control in the primary visual cortex but rather gating in parietal regions (Zhigalov & Jensen, 2020). With our new insight, spatial attention paradigms could be used to uncover the visuo-parietal information flow by assessing the phase relationship between the primary visual and parietal cortices.

### Functional role of parietal regions

In our recent study (Zhigalov & Jensen, 2020), we found that spatial attention modulates both frequency tagging response in the early visual cortex and the alpha band oscillations in parietal regions. Importantly the alpha activity and the frequency tagging response were not correlated over trials. This pointed to a scheme in which alpha oscillations gate the information flow in downstream visual regions without directly controlling the gain in the early visual cortex. In a similar study (Sokoliuk et al., 2019), two sources of alpha oscillations in the primary visual cortex and parietal areas were identified. These sources were both modulated by attention but played different functional roles depending on behavioural demands. The stage is now set for investigating how the visual and parietal interactions are affected by the difference in alpha phase between the two generators in relation to cognitive demands.

## Acknowledgement

The work was supported by the following funding: a James S. McDonnell Foundation Understanding Human Cognition Collaborative Award (grant number 220020448), the Wellcome Trust Investigator Award in Science (grant number 207550), a BBSRC grant (BB/R018723/1) as well as the Royal Society Wolfson Research Merit Award. AZ acknowledges financial support from the University of Birmingham Dynamic Investment Fund.

## Notes

### Competing Interest Statement

The authors have declared no competing interest.

### Summary of Updates

Minor corrections in the text

